# Assessing the impact of static and fluctuating ocean acidification on the behavior of *Amphiprion percula*

**DOI:** 10.1101/2021.01.23.427511

**Authors:** Matthew A. Vaughan, Danielle L. Dixson

## Abstract

Coral reef organisms are exposed to both an increasing magnitude of *p*CO_2_, and natural fluctuations on a diel scale. For coral reef fishes, one of the most profound effects of ocean acidification is the impact on ecologically important behaviors. Previous behavioral research has primarily been conducted under static *p*CO_2_ conditions and have recently come under criticism. Recent studies have provided evidence that the negative impacts on behavior may be reduced under more environmentally realistic, fluctuating conditions. We investigated the impact of both present and future day, static (500 and 1000 μatm) and diel fluctuating (500 ± 200 and 1000 ± 200 μatm) *p*CO_2_ on the lateralization and chemosensory behavior of juvenile anemonefish, *Amphiprion percula*. Our static experimental comparisons support previous findings that under elevated *p*CO_2_, fish become un-lateralized and lose the ability to discriminate olfactory cues. Diel-fluctuating *p*CO_2_ may aid in mitigating the severity of some behavioral abnormalities such as the chemosensory response, where a preference for predator cues was significantly reduced under a future diel-fluctuating *p*CO_2_ regime. This research aids in ground truthing earlier findings and contributes to our growing knowledge of the role of fluctuating conditions.

## 1. Introduction

Anthropogenic climate change is rapidly altering Earth’s oceans, leading to the phenomenon of ocean acidification (OA). The burning of fossil fuels from human activities is exhausting excessive concentrations of carbon dioxide (CO_2_) into the atmosphere at an unprecedented rate (Kerr 2010). The ocean sequesters atmospheric CO_2_ by uptaking over one third of anthropogenic CO_2_ emissions (Sabine et al. 2004; Rhein et al. 2013). The additional input of atmospheric CO_2_ reacts with seawater to acidify the oceans (Hoegh-Guldberg et al. 2007; Doney et al. 2009).

From the pre-industrial era, global ocean pH has decreased by at least 0.1 units (Wooton et al. 2008). Under “business as usual” conditions, it is forecasted that ocean pH will decrease by up to 0.43 units by the end of the century (RCP8.5 scenario; IPCC 2014). If these future projections hold true, changes in ocean carbonate chemistry will be unlike any past event (Hönisch et al. 2012).

Ocean acidification has been shown to affect the sensory system, physiology, and behavior of marine fishes (Heuer and Grosell 2014; Ashur et al. 2017; Cattano et al. 2018). Some of the most notable effects of OA on coral reef fishes are on ecologically important behaviors (Nagelkerken and Munday 2016). These behavioral changes include disruption to olfactory preferences (Munday et al. 2009; Dixson et al. 2010; Ferrari et al. 2012a), reduced prey detection (Cripps et al. 2011), decreased learning ability (Ferrari et al. 2012b), reduced behavioral lateralization (Domenici et al. 2012, 2014; Nilsson et al. 2012), increased activity and boldness (Munday et al. 2010, 2014; Nilsson et al. 2012; Nagelkerken and Munday 2016), reduced hearing and vision (Simpson et al. 2011; Chung et al. 2014), and altered reaction times, escape speeds and distances (Allan et al. 2013, 2017; Munday et al. 2016). Furthermore, elevated CO_2_ exposure can affect coral reef fish settlement behavior (Devine et al. 2012), shoaling behavior (Nadler et al. 2016), habitat preference (Devine and Munday 2013; Nagelkerken and Munday 2016; Goldenburg et al. 2018), and the replenishment of fish stocks (Munday et al. 2010).

The effect of OA on coral reef fish juveniles is not ubiquitous across all species, even within the same genus, with some being more tolerant to increased CO_2_ than others (Ferrari et al. 2011; McCormick et al. 2013). An analysis of the OA literature shows large variation in the sensitivity of behavioral responses to OA, at habitat and environmental scales, as well as interspecific and intraspecific scales (McCormick et al. 2013; Clements and Hunt 2015; Schunter et al. 2016; Vargas et al. 2017; Cattano et al. 2018; Munday et al. 2019, Munday et al. 2020).

The impacts of future climate change scenarios on coral reef fish behavior are not yet completely understood. Stressors such as OA occur under fluctuating regimes and in synergy with other stressors. Marine organisms experience a range of pH variability on both temporal and spatial scales dependent on their environment, and these fluctuations are expected to be exacerbated under future climate change conditions (Hofmann et al. 2011; Johnson et al. 2013; Shaw et al. 2013; Kapsenberg et al. 2015; McNeil and Matsumoto 2019). Coral reefs experience diel *p*CO_2_ fluctuations due to a range of processes including biological reef metabolism (Waldbusser and Salisbury 2014), as well as seasonal variability, ranging greater than 200 μatm at times (Shamberger et al. 2011; Price et al. 2012; Albright et al. 2013; Duarte et al. 2013; Kline et al. 2015). Furthermore, fish seldom remain in one location and will likely experience a wide range of pH variability over their life history due to movement around and off the reef, and ecosystem structure and function can vary across different locations on reefs due to variability in pH (Price et al. 2012; Shaw et al. 2013). There is evidence to suggest that temporary movement to another water source (e.g. from an area of low pH to an area of higher pH) will not immediately influence the behavioral changes resulting from elevated CO_2_ (Munday et al. 2016).

The majority of previous studies have assessed behavioral changes using static OA treatment conditions, however, considering the importance of better reflecting the natural environment, a greater understanding of the impacts fluctuating conditions have on behavior is required. Little research has been conducted on the impact fluctuating stressors may have on the behavior of coral reef fish. Recent studies have shown that diel pH fluctuations may offset or reduce the severity of behavioral changes when compared to static future OA conditions (Ou et al. 2015; Jarrold et al. 2017; Jarrold and Munday 2018) suggesting that the findings of earlier research (conducted under static OA conditions) may have overestimated the degree of behavioral impairment.

Key behavioral traits exhibited by coral reef fishes such as behavioral lateralization and chemosensory response have primarily been studied under static OA conditions (Munday et al. 2009, 2010, 2013; Dixson et al. 2010; Cripps et al. 2011; Ferrari et al. 2011, 2012a, 2012b; Devine et al. 2012; Domenici et al. 2012, 2014; Nilsson et al. 2012; Devine and Munday 2013; Welch et al. 2014). These behaviors play important ecological roles and are often crucial for survival. The inability to generalize OA findings has resulted in a challenge to the understanding of the behavioral impacts of OA on coral reef fishes, with differing results collected on alternative species, life history stages, methods and testing apparatus being used (Munday et al. 2020). These discrepancies make it important to revisit and accurately replicate previous studies to determine if earlier findings hold true, while still advancing the research field through the inclusion of diel fluctuations. Technological advancements have now made it possible to test these behavioral traits under more realistic and biologically relevant environmental conditions. It is largely unknown if, and to what degree, the brief release from low pH in the evenings will offset the negative effects of OA. The objectives of this research were to assess the impacts future climate change conditions will have on the behavior of juvenile coral reef fish, *Amphiprion percula*, under both static conditions (to revisit and replicate previous findings) and under more ecologically relevant conditions (i.e. fluctuating conditions). Two experimental trials were conducted investigating: 1) behavioral lateralization; 2) the chemosensory response.

## 2. Materials and Methods

### 2.1. Study Species

A total of 169 chemically naïve, laboratory bred juvenile *Amphiprion percula* were sourced from Sustainable Aquatics (Jefferson City, TN). All fish were bred from wild stock parents that were randomized to include the offspring of three parental groups to account for any genetic differences. Fish were 18 weeks old at the commencement of the experiment. Juvenile *A. percula* were fed 0.8 mm pellets (Sustainable Aquatics) and *Artemia* sp. nauplii daily in the morning of the first two weeks, before transitioning to a pellet-only diet during behavioral trials. During behavioral trials, fish were fed at the end of the trial period on test days. Fish were maintained on a 12:12 hr light:dark cycle.

### 2.2. Experimental Design and Protocol

The impact of static and fluctuating OA on behavioral changes was assessed using current day static temperature (28.5 °C) with a 2 × 2 cross-factorial design. Treatment groups included: 1) static present-day control conditions; 2) static future-day acidification conditions; 3) fluctuating present-day control conditions; and 4) fluctuating future-day acidification conditions. Each treatment group consisted of 5 replicate 20 L aquariums (26 cm × 26 cm × 31 cm) holding 7-9 *A. percula*. Juveniles were habituated to their treatment conditions for 15 days prior to commencement of behavioral trials. A habituation timeframe of 4-7 days has proven to be sufficient to impair a range of behavioral responses (Munday et al. 2010; Devine and Munday 2013; Chivers et al. 2014).

Aquaria were separated into one of two large re-circulating systems (111 cm × 248 cm × 22 cm), each holding two treatment groups in a water bath. Natural seawater was used and sourced from the Indian River Inlet (Delaware, USA). A header sump (680 L) pumped water into each of the aquaria at an adjusted flow rate (75 mL min^-1^). Four APEX Fusion systems (Neptune Systems) were used to independently control temperature and pH_NBS_ based on programmed set points. CO_2_ was regulated and monitored using an APEX computer system. This system injects additional CO_2_ when pH levels exceed the desired computer set point by opening a solenoid connected to a precision needle valve. A steady slow stream of CO_2_ was bubbled into the tank through an air stone until the pH has dropped below the set maximum. CO_2_-stripped air (achieved using a soda lime filter) was also bubbled into each aquarium system at a controlled flow rate to raise and maintain pH at desired values. Both control and future day treatments used the same methods, however pH set points for each varied. Temperature was monitored at 2 min intervals using Neptune temperature probes controlled by the APEX system and adjusted with 200 W heaters (ViaAqua) placed in each tank. When the temperature probe read below the desired set point, power was turned on to the tank’s heater, and the heater was then turned off when the temperature reached the set point.

Both pH and temperature were independently tested twice daily using a handheld Mettler Toledo probe (SevenGo Duo pro SG68 pH meter). Salinity was tested twice daily using a handheld refractometer (Fisherbrand Salinity Refractometer). Partial water changes were performed every second day to control for salinity. Water quality testing (ammonia, nitrate, nitrite) was conducted twice weekly.

### 2.3. Carbonate Chemistry

Juvenile *A. percula* were treated for 15 days (4^th^ – 19^th^ Nov 2019) in either static present day (500 μatm), static future day (1000 μatm), fluctuating present day (1000 ± 200 μatm), or fluctuating future day (500 ± 200 μatm) CO_2_ treatments at a current day static temperature of 28.5 °C (Table 1). Treatment continued throughout the next 15 days during the behavioral trial period (20^th^ Nov – 4^th^ Dec 2019). The control pH levels were based on the averages of five present-day reef systems (Fig. 1; Hofmann et al. 2011; Albright et al. 2013). The targeted static future day treatment values (~1000 μatm) were based on commonly used forecast open ocean values for the end of the century (Kroeker et al. 2013; IPCC 2014). Coral reef *p*CO_2_ fluctuations can typically range between ±50-150 μatm on a diel scale (Albright et al. 2013; Kline et al. 2015). The magnitude in range of fluctuations is forecast to increase in the future (McNeil and Sasse 2016), and this experiment aimed for a range of ±200 μatm, as seen in Jarrold at el. (2017). Fluctuating treatment groups had hourly shifts in pH_NBS_ of 0.01 – 0.02 units to reflect diel fluctuations that occur on a natural reef (Fig. 1). Future day treatment levels used the control levels as a baseline and were adjusted to include the forecast values expected to occur in 2100 (IPCC 2014), where fluctuations were extrapolated to 0.3 units lower than control levels. The pH_NBS_ set points were determined through prior experimentation and programmed into the APEX system to correspond with target *p*CO_2_ values. Temperature levels were based on summer values recorded by Albright et al. (2013). Throughout the experimental period mean temperature was 28.54 ± 0.02 °C, and mean salinity was 34.43 ± 0.06 ppm. To determine the carbonate chemistry of each treatment group, dissolved inorganic carbon (DIC) and pH samples were taken each week. The certified reference materials provided by the laboratory of Dr. A Dickson (San Diego, CA, USA) with known DIC were used to validate DIC measurements, and pH was measured spectrophotometrically (Dickson et al. 2007). DIC and spectrophotometric pH were used to calculate*pCO_2_* with the CO2SYS software (Pierrot et al. 2006) using constants K1 and K2 (Mehrbach et al. 1973) and refit by Dickson and Millero (1987). Measurements for the fluctuating treatments were taken midweek at multiple times to correspond with the highest, median, and lowest expected *p*CO_2_ values (Table 1.).

**Table 1.**
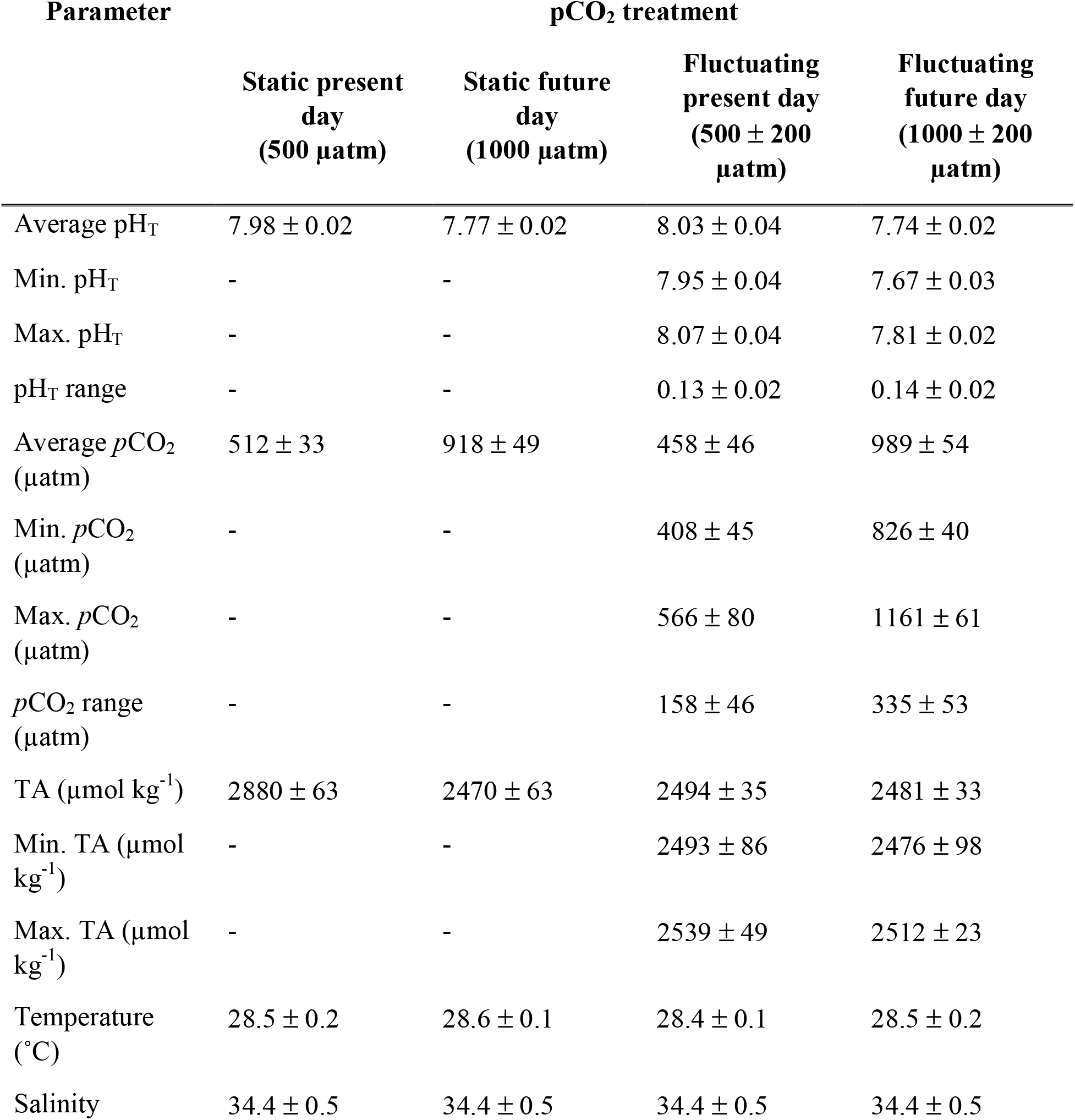
Mean values for seawater parameters ± 1 SD throughout the entirety of the experimental duration. Average values of *p*CO_2_ and pH_T_ for fluctuating treatments are from samples taken at different times of the day to reflect the middle, minimum, and maximum values. Range of *p*CO_2_ and pH_T_ represent the average range of all replicates within a treatment between the minimum and maximum values. Temperature values provided are from a portable Mettler Toledo probe.

| Parameter | pCO<sub>2</sub> treatment |  |  |  |
| --- | --- | --- | --- | --- |
| | Static present day<br>(500 $\mu\text{atm}$) | Static future day<br>(1000 $\mu\text{atm}$) | Fluctuating present day<br>(500 ± 200 $\mu\text{atm}$) | Fluctuating future day<br>(1000 ± 200 $\mu\text{atm}$) |
| Average $\text{pH}_T$ | 7.98 ± 0.02 | 7.77 ± 0.02 | 8.03 ± 0.04 | 7.74 ± 0.02 |
| Min. $\text{pH}_T$ | - | - | 7.95 ± 0.04 | 7.67 ± 0.03 |
| Max. $\text{pH}_T$ | - | - | 8.07 ± 0.04 | 7.81 ± 0.02 |
| $\text{pH}_T$ range | - | - | 0.13 ± 0.02 | 0.14 ± 0.02 |
| Average $p\text{CO}_2$<br>($\mu\text{atm}$) | 512 ± 33 | 918 ± 49 | 458 ± 46 | 989 ± 54 |
| Min. $p\text{CO}_2$<br>($\mu\text{atm}$) | - | - | 408 ± 45 | 826 ± 40 |
| Max. $p\text{CO}_2$<br>($\mu\text{atm}$) | - | - | 566 ± 80 | 1161 ± 61 |
| $p\text{CO}_2$ range<br>($\mu\text{atm}$) | - | - | 158 ± 46 | 335 ± 53 |
| TA ($\mu\text{mol kg}^{-1}$) | 2880 ± 63 | 2470 ± 63 | 2494 ± 35 | 2481 ± 33 |
| Min. TA ($\mu\text{mol kg}^{-1}$) | - | - | 2493 ± 86 | 2476 ± 98 |
| Max. TA ($\mu\text{mol kg}^{-1}$) | - | - | 2539 ± 49 | 2512 ± 23 |
| Temperature<br>($^{\circ}\text{C}$) | 28.5 ± 0.2 | 28.6 ± 0.1 | 28.4 ± 0.1 | 28.5 ± 0.2 |
| Salinity | 34.4 ± 0.5 | 34.4 ± 0.5 | 34.4 ± 0.5 | 34.4 ± 0.5 |

**Fig 1.**
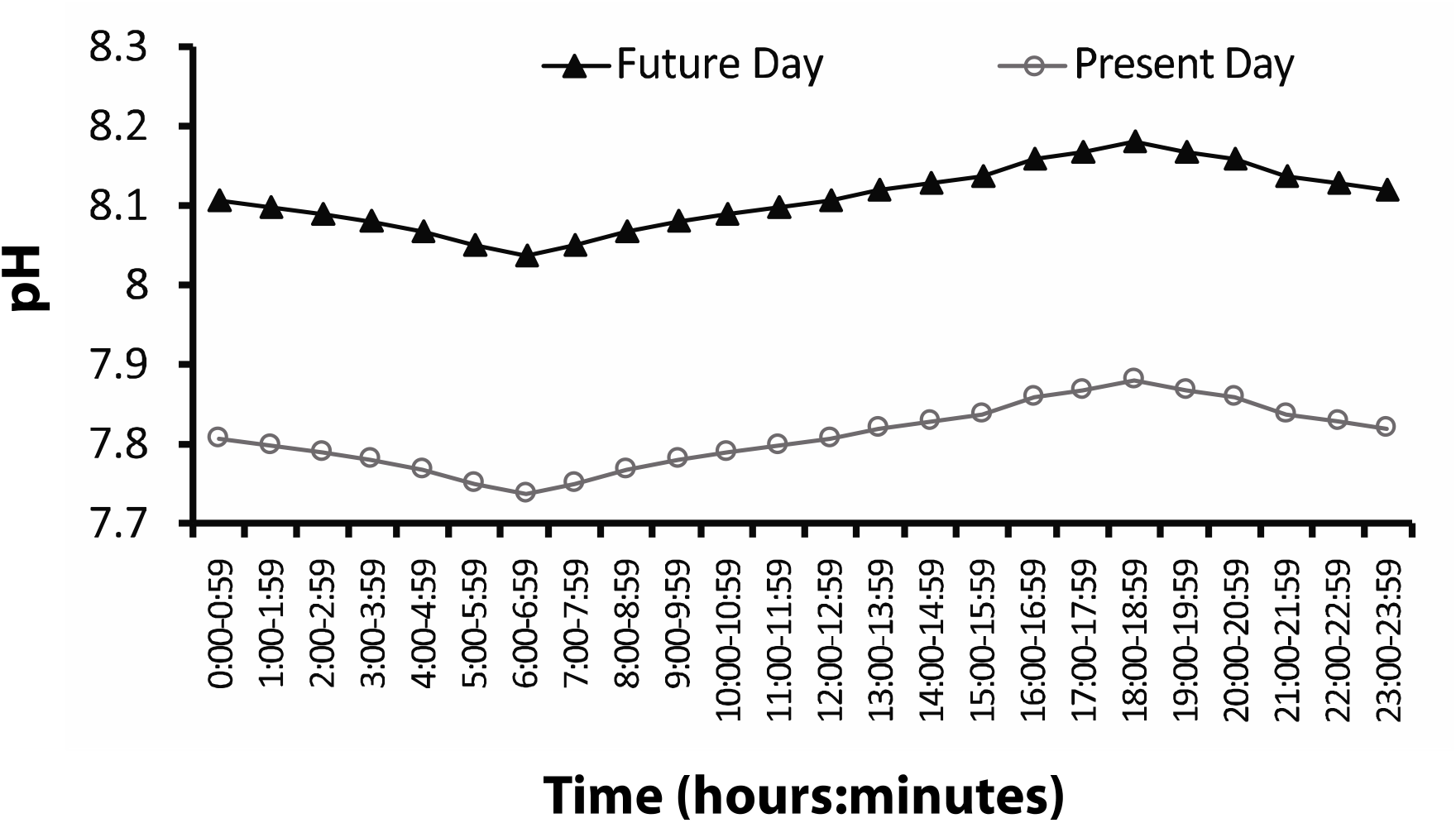
Target diel pH fluctuations for present day control (black, closed triangles) and future day treatment (gray, opened circles) tanks. APEX probe error ± 0.02.

### 2.4. Behavioral Trials

Two experimental trials measuring ecologically relevant behaviors were conducted investigating 1) behavioral lateralization, and 2) chemosensory response. Trials were run sequentially, and each trial was conducted once from a haphazardly selected sample of juvenile fish from each treatment group. Each individual fish was only tested once per experimental trial. All behavioral trials were undertaken between 08:00 – 17:00. Additional sensory stimuli were minimized throughout all trials.

Behavioral lateralization, which reflects the tendency for fish to have a left or right turning preference, was evaluated using a detour test following similar methods to Domenici et al. (2012). A two-way T-maze was used to measure the turning direction of an individual. Water from the respective treatment conditions of each fish was used to fill the maze to a depth of 4 cm. A single fish was placed at one end of the T-maze where it could explore the maze and habituate for 3 min (n=30). Following the habituation period, the fish was gently coaxed with a plastic rod (no closer than two body lengths away from the fish) to the middle of the runway, then through the maze to the end of the runway where it was confronted with a turning choice of either left or right. Ten consecutive runs were recorded for each fish, and the score of the turning direction and the degree of lateralization was obtained. Direction choice was determined as the first direction chosen when the fish exited the runway. The observer was kept blind to the treatment during the trial.

An Atema two-channel choice flume (Atema et al. 2002) was used to test the chemosensory mediated behavior towards predator chemical signals by juvenile *A. percula*, following the methods of Gerlach et al. (2007). Chemical cues for testing were generated by soaking either a single predator (*Cephalopholis cyanostigma*) or a single non-predator (*Zebrasoma falvescens*) in a closed, aerated 10 L seawater system for 2 hr. The seawater used was from the same source as the seawater in the fish aquaria, which had not been treated during the soak period. The predator or non-predator was then removed from the water immediately following the soak. Cue water was then placed in the system to treat the water to the relevant temperature and pH associated with the fish being tested. This process ranged from 10 – 30 min depending on the treatment conditions. To run the two-channel choice flume, water from two sources (either predator, non-predator, or untreated control water) were gravity fed into the flume (13 cm × 4 cm) at 100 mL min^-1^. Water velocity was controlled using two flow meters, ensuring water was delivered at equal rates to prevent mixing of the water masses and to achieve laminar flow for the cues. Each individual fish was isolated in a small 200 mL beaker of associated treatment water for 10 min, and then gently placed downstream in the flume and given a 2 min habituation period where it was free to swim throughout the chamber. At the conclusion of the habituation period, the fish’s position on either the right or left side was recorded at 5-second intervals for 2 min. The water sources were then switched to discount a side preference, and the flume was then given a 1 min flushing period before the entire 2 min habituation period and 2 min testing period were repeated. Dye tests were conducted before and after trials and regularly throughout to ensure laminar flow with no areas of turbulence or eddies. Juvenile fish were haphazardly selected from each treatment group (n=20). Three runs were conducted in the same order for each fish: 1) non-predator vs. untreated, 2) predator vs. untreated, 3) non-predator vs. predator. All trials were run double blinded, with both fish treatment group and chemical cues blind to the observer.

### 2.5. Statistical Analysis

To assess both the turning preference of the fish and the strength of lateralization, relative lateralization (*L*_R_) and absolute lateralization (*L*_A_) were calculated using established methods (Bisazza et al. 1998). To compare differences between treatment groups, a Kruskal-Wallis test was performed on both relative and absolute lateralization, as the static present day data set was not normal (D’Agostino & Pearson omnibus normality test, p = 0.0336). Additionally, lateralization was also assessed at both the population-level using a generalized linear random-effects model (GLMM; using the lme4 package in R), and at the individual-level using a chisquare test, following previously used methods (Roche et al. 2020). Shapiro-Wilk normality tests were performed on the chemosensory response data. To assess if there was a significant preference towards a specific cue within a treatment group per chemosensory response trial, one sample *t*-tests were conducted comparing the mean and variance against an expected value of 50%. To compare if there were differences between treatment groups, data was first transformed using the arcsine (square root) transformation. A one-way ANOVA with Tukey’s posthoc test was run on each of the three different chemosensory trials.

## 3. Results

### 3.1. Behavioral lateralization

All treatment groups exhibited a mean left turning preference (Fig. 2A), where fish from the static future day (SFD) and fluctuating future day (FFD) treatments displayed a higher degree of mean relative lateralization (*L*_R_) (SFD: −9.33 ± 6.49, FFD: −6.67 ± 6.80). However, there were no significant differences between treatments for *L*_R_ (Kruskal-Wallis, p = 0.9254), and no treatment group was lateralized at the population level (GLMM, p >0.05 for all cases).

**Fig 2.**
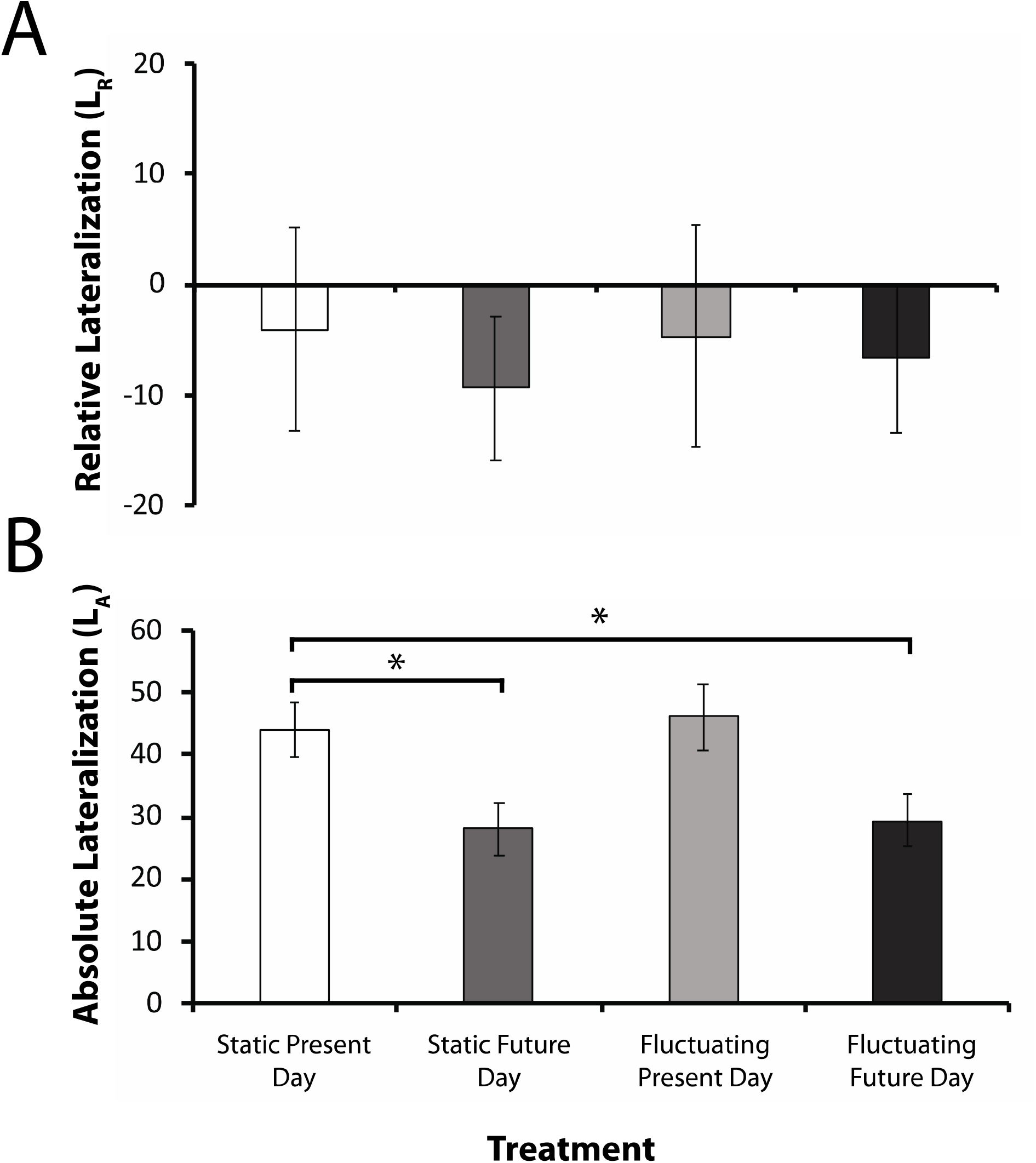
**A)** Relative lateralization (*L*_R_) of juvenile *Amphiprion percula* (mean ± S.E.). Positive and negative values on the y-axis indicate either a right or left group turning preference, respectively. **B)** Absolute lateralization (*L*_A_) of juvenile *Amphiprion percula* (mean ± S.E.). Significant differences (p < 0.05) between groups are represented by an asterisk. A total of 30 fish from each *p*CO_2_ treatment group were used in the detour test.

There were significant differences between treatments for absolute lateralization (*L*_A_) (Kruskal-Wallis, p <0.05). Fish from the static present day (SPD) and fluctuating present day (FPD) controls exhibited the highest mean values of *L*_A_ (SPD: 44 ± 4.54, FPD: 46 ± 5.27) (Fig. 2B). In contrast, fish from the static future day and fluctuating future day treatments displayed lower mean *L*_A_ values (SFD: 28 ± 4.25, FFD: 29.33 ± 4.26). Fish from both present day controls remained individually lateralized (χ^2^, SPD: p <0.00001, FPD: p <0.000001), whereas fish from future day treatments lost their individual lateralization (χ^2^, SFD: p = 0.1571; FFD: p = 0.0797). Fish treated with static and fluctuating future day conditions displayed significant differences in *L*_A_ with fish from the static present day control (Kruskal-Wallis, p <0.05 in both cases), but not with fish from the fluctuating present day control (Kruskal-Wallis, SFD: p = 0.0673; FFD: p = 0.0544). There were no significant differences in *L*_A_ between both static and fluctuating present day (Kruskal-Wallis, p = 0.8624) and static and fluctuating future day (Kruskal-Wallis, p = 1) treatments.

### 3.2. Chemosensory response

Fish treated with static present day, fluctuating present day, and fluctuating future day seawater displayed no preference for the chemical cues produced by the non-predator (*Zebrasoma falvescens*) when tested against untreated seawater (*t*-test, p >0.05), spending between 45.4 – 54.3% (SPD: 54.3 ± 2.2%, FPD: 45.4 ± 2.1%, FFD: 49.2 ± 5.0%) time in the non-predator chemical cues (Fig. 3A; Table 2.). Fish in the static future day treatment group showed a significant preference toward the untreated control water (SFD: 54.1 ± 3.2%; *t*-test, p = 0.042). There were no significant differences between treatment groups for time spent in cue (ANOVA, p >0.05; Table 3.).

**Fig 3.**
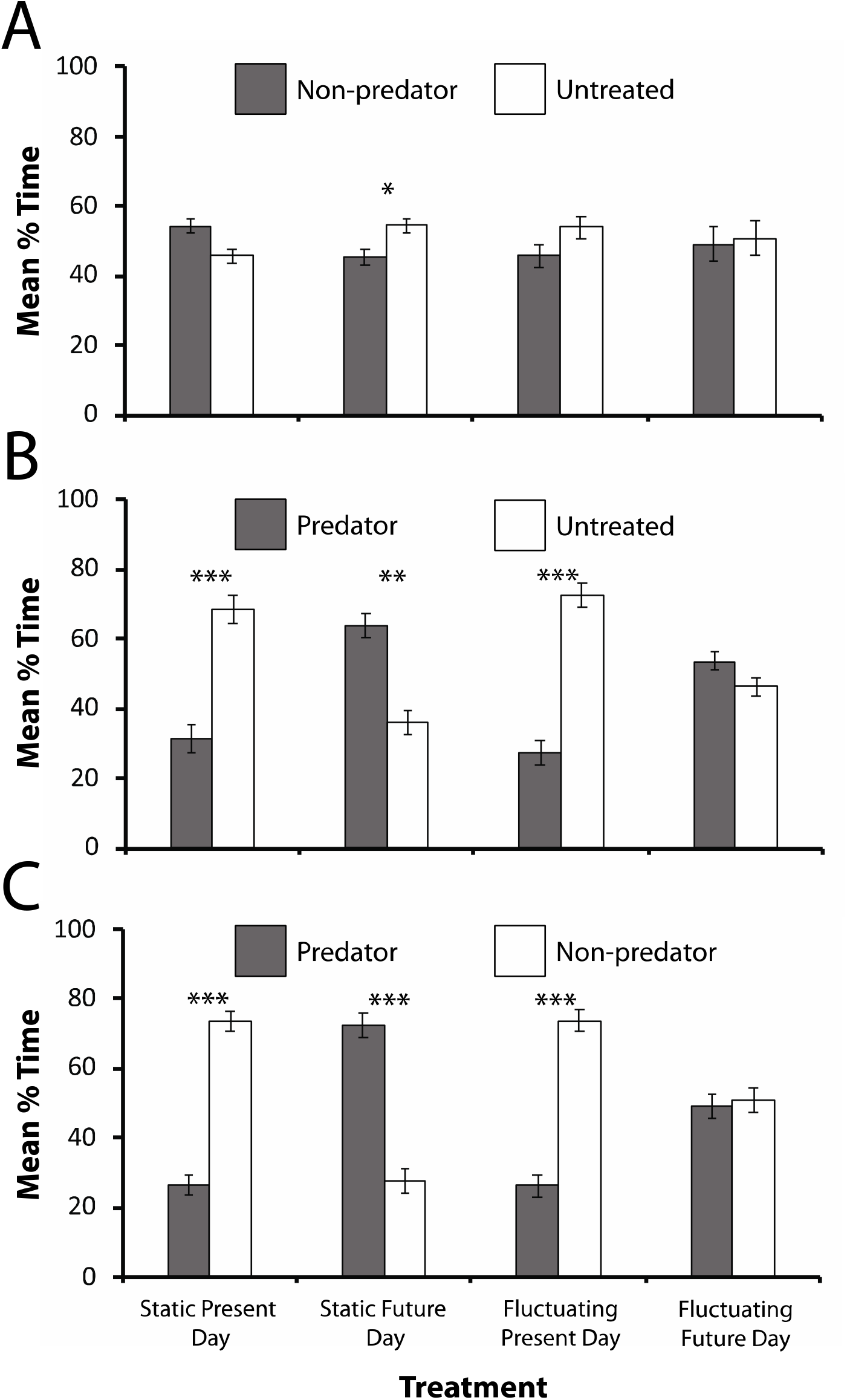
Mean percentage of time (± S.E.) juvenile *Amphiprion percula* spent in different chemical cues presented in a two-channel choice flume. A total of 20 fish from each *p*CO_2_ treatment group were used per chemical cue trial with a choice of: **A**) non-predator (Tang) and untreated (control), **B**) predator (Cod) and untreated (control), and **C**) predator (Cod) and non-predator (Tang). Significant differences between chemical cue preference within treatment groups are represented by asterisks, where: * = p < 0.05; ** = p < 0.001; *** = p <0.00001.

**Table 2.** Comparison of percent time (± SE) fish from each treatment group spent in either chemical cue presented in each of the three chemosensory trials (n=30). The p-values represent one-sample *t*-tests conducted on percent time spent in cue against an expected value of 50%. Significance values are reflected with an asterisk and represent a preference toward the cue tested (left side chemical cue comparison column).

| Treatment | Fluctuation | Chemical cue comparison (%) |  | p-value |
| --- | --- | --- | --- | --- |
|  |  | Non-predator | Untreated seawater |  |
| Present Day | Static | 54.27 ± 2.16 | 45.73 ± 2.16 | 0.062 |
| Present Day | Fluctuating | 45.94 ± 3.22 | 54.06 ± 3.22 | 0.222 |
| Future Day | Static | 45.42 ± 2.10 | 54.58 ± 2.10 | 0.042 * |
| Future Day | Fluctuating | 49.17 ± 4.95 | 50.83 ± 4.95 | 0.860 |
|  |  | Predator | Untreated seawater |  |
| Present Day | Static | 31.46 ± 4.29 | 68.54 ± 4.29 | $3.702 \times 10^{-4}$ * |
| Present Day | Fluctuating | 63.85 ± 3.33 | 36.15 ± 3.33 | $5.849 \times 10^{-6}$ * |
| Future Day | Static | 27.40 ± 3.64 | 72.60 ± 3.64 | $5.409 \times 10^{-4}$ * |
| Future Day | Fluctuating | 53.65 ± 2.48 | 46.35 ± 2.48 | 0.158 |
|  |  | Predator | Non-predator |  |
| Present Day | Static | 26.46 ± 2.78 | 73.54 ± 2.78 | $6.935 \times 10^{-8}$ * |
| Present Day | Fluctuating | 72.50 ± 3.41 | 27.50 ± 3.41 | $6.752 \times 10^{-7}$ * |
| Future Day | Static | 26.35 ± 3.25 | 73.65 ± 3.25 | $2.6 \times 10^{-6}$ * |
| Future Day | Fluctuating | 49.06 ± 3.57 | 50.94 ± 3.57 | 0.611 |

**Table 3.** Comparison of percent time (± SE) fish from each treatment group spent in either chemical cue presented in each of the three chemosensory trials (n=30). The p-values represent comparisons between treatment groups in each chemical cue (conducted via ANOVA), where: SPD = Static Present Day; SFD = Static Future Day; FPD = Fluctuating Present Day; and FFD = Fluctuating Future Day. Asterisks reflect significant differences.

| Treatment | Fluctuation | Chemical cue comparison (%) |  | p-value |
| --- | --- | --- | --- | --- |
|  |  | Non-predator | Untreated seawater |  |
| Present Day | Static | 54.27 ± 2.16 | 45.73 ± 2.16 | SPD-SFD: 0.493<br>SPD-FPD: 0.417<br>SPD-FFD: 0.736 |
| Future Day | Static | 45.94 ± 3.22 | 54.06 ± 3.22 | SFD-FPD: 0.999<br>SFD-FFD: 0.979 |
| Present Day | Fluctuating | 45.42 ± 2.10 | 54.58 ± 2.10 | FPD-FFD: 0.954 |
| Future Day | Fluctuating | 49.17 ± 4.95 | 50.83 ± 4.95 | |
|  |  | Predator | Untreated seawater |  |
| Present Day | Static | 31.46 ± 4.29 | 68.54 ± 4.29 | SPD-SFD: $1.5 \times 10^{-6}$ *<br>SPD-FPD: 0.888<br>SPD-FFD: $8.359 \times 10^{-4}$ * |
| Future Day | Static | 63.85 ± 3.33 | 36.15 ± 3.33 | SFD-FPD: $1 \times 10^{-7}$ *<br>SFD-FFD: 0.353 |
| Present Day | Fluctuating | 27.40 ± 3.64 | 72.60 ± 3.64 | FPD-FFD: $6.21 \times 10^{-5}$ * |
| Future Day | Fluctuating | 53.65 ± 2.48 | 46.35 ± 2.48 | |
|  |  | Predator | Non-predator |  |
| Present Day | Static | 26.46 ± 2.78 | 73.54 ± 2.78 | SPD-SFD: $<1 \times 10^{-7}$ *<br>SPD-FPD: 0.999<br>SPD-FFD: 0.003 * |
| Future Day | Static | 72.50 ± 3.41 | 27.50 ± 3.41 | SFD-FPD: $<1 \times 10^{-7}$ *<br>SFD-FFD: $1.296 \times 10^{-4}$ * |
| Present Day | Fluctuating | 26.35 ± 3.25 | 73.65 ± 3.25 | FPD-FFD: 0.004 * |
| Future Day | Fluctuating | 49.06 ± 3.57 | 50.94 ± 3.57 | |

When the predator chemical cue (*Cephalopholis cyanostigma*) was tested against untreated seawater, fish held in future day conditions displayed a preference for the predator chemical cues, spending between 53.7 – 63.9% (SFD: 63.9 ± 3.3%, FFD: 53.7 ± 2.5%) of their time in the predator chemical cue (Fig. 3B; Table 2.). A statistically significant preference between predator cue and control water was identified for the static future day treatment (*t*-test, p <0.001) but not for the fluctuating future day treatment (*t*-test, p = 0.158). Significant differences in the time spent in predator cue between treatments were identified (ANOVA, p <1 × 10^-8^; Table 3). The preferences observed in the future day treatments significantly differed in comparison to present day treatments groups, where fish from the static future day treatment spent significantly more time in the predator cue compared to the static (ANOVA, p <1 × 10^-5^) and fluctuating (ANOVA, p <1 × 10^-6^) present day treatments. The same was true for fish in the fluctuating future day treatment compared to the static (ANOVA, p <0.001) and fluctuating (ANOVA, p <1 × 10^-4^) present day treatment groups. There was no significant difference in time spent in predator cue between fish from the fluctuating future day treatment and static future day treatment (ANOVA, p >0.05; Fig. 3B).Fish from both present day treatments (static and fluctuating) spent significantly more time in the untreated water over the predator cue (*t*-test, SPD: p <0.001; FPD: p <1 × 10^-6^), spending 68.5 – 72.6% of their time in the untreated water (SPD: 68.5 ± 4.3%, FPD: 72.6 ± 3.6%). No significant difference in percent time spent in the predator cue between present day treatments was found (ANOVA, p = 0.888).

When comparing the predator chemical cues to the non-predator chemical cues simultaneously (i.e. *C. cyanostigma* vs. *A. pyropherus*), a significant preference for the predator chemical cue was displayed by fish treated in static future day conditions (*t*-test, p <1 × 10^-5^; Fig. 3C), spending almost three times longer in this cue. The CO_2_ treatment of the fish resulted in significantly different responses to the predator cue (ANOVA, p <1 × 10^-11^; Table 3.). Fish from the static future day treatment spent significantly more time in the predator cue than all other treatments (ANOVA, SPD: p <1 x 10^-7^; FPD: p <1 × 10^-7^; FFD: p <0.001). Fish treated under fluctuating future day conditions displayed no significant preferences for the predator or non-predator (*t*-test, p = 0.6111). This was a significantly greater amount of time in comparison to the static and fluctuating present day treatments (ANOVA, p <0.01 in both cases), but notably significantly less time in comparison to the static future day treatment (ANOVA, p <0.001; Fig. 3C). Fish from both present day treatments spent a significant amount of time in the chemical cues produced by the non-predator over the predator (*t*-test, SPD: p <1 × 10^-7^; FPD: p <1 × 10^-6^), but this did not significantly differ between the two groups (ANOVA, p = 0.9999).

## 4. Discussion

This study supports most previous research assessing the impact of ocean acidification on juvenile coral reef fish behavior. In both behavioral trials conducted, fish in the future day treatments exhibited behavioral changes that are likely deleterious in comparison to fish in present day treatments. When provided with the option of a predator cue compared to either untreated control water or non-predator cue, fish from the static future day treatment spent significantly more time in the predator cue compared to fish treated with present day conditions. Diel *p*CO_2_ fluctuations had varying impacts on juvenile *A. percula* behavior depending on whether fish were treated under present day or future day conditions. When comparing fluctuating conditions with static conditions for fish treated in future day *p*CO_2_, fluctuations reduced the attraction (i.e. amount of time) toward the predator cue. Fish from the fluctuating future day treatment had no significant preference between the non-predator and predator cues, where under static future day conditions they preferred the predator cue, thus indicating that fluctuating conditions may help mitigate the degree of negative behavioral impairment on chemosensory response. In contrast, fluctuations did not reduce the amount of time fish from present day treatments spent in the predator cue. These results suggest that natural diel fluctuations may help mitigate negative behavioral impairment in the future, but under present day conditions there is no apparent influence on behaviors tested in this study.

The results from our behavioral lateralization trials support those of previous findings, where fish from both future day treatments (static and fluctuating) became un-lateralized, while fish from present day conditions remained individually lateralized (Domenici et al. 2012, 2014). However, fluctuating conditions did not appear to mitigate or offset this behavioral change, suggesting fluctuating conditions may not aid in behavioral lateralization abnormalities. Contrasting this, Jarrold et al. (2017) found that fluctuating future day conditions did reduce the negative behavioral impairment of *A. percula* becoming un-lateralized in their study. Given the small number of studies conducted in this field at the time of writing, more research is required to garner a better picture of how behavioral lateralization may be impacted under more realistic, future day conditions. Fish from all treatment groups displayed a left turning preference, differing from previous findings of other juvenile coral reef damselfish that have showed varied results (Domenici et al. 2012, 2014; Jarrold et al. 2017). Recent research has suggested that behavioral lateralization, through the methods of a detour test, is not repeatable in fishes (Roche et al. 2020). When five different species of fish were run multiple times, there was no repeatability in results. Although we did not repeat behavioral lateralization trials multiple times, given the high degree of both interspecific and intraspecific sensitivity to OA, this suggestion may not hold true to all fishes. For example, a recent study found that behavioral lateralization is repeatable across contexts (McLean and Morell 2020). In this study, relative lateralization was repeatable for male and female adult *Poecilia reticulata*, and absolute lateralization was repeatable for males. It is also possible that fish could learn the detour maze or become unthreatened when placed in the same situation repeatedly. Future day *p*CO_2_ conditions impacted the ability of juvenile coral reef fish to discriminate between different cue sources, as fish from the present day treatments showed an attraction toward the non-predator cue when given the option between a predator and non-predator cue. A higher percent of time spent in the predator cue from fish treated with future day *p*CO_2_ conditions is likely due to a loss of discriminatory ability rather than an attraction toward the predator cue, supporting previously reported findings of attraction toward unfamiliar settlement cues and predator cues (Munday et al. 2009, 2016; Dixson et al. 2010; Nilsson et al. 2012).

Changes in chemosensory discrimination may also result in higher mortality rates (Munday et al. 2014). However, our results indicate that naturally occurring diel *p*CO_2_ fluctuations may, to a degree, mitigate the impact of these negative behavioral impairments. Jarrold et al. (2017) found similar reductions in time spent in predator cues under fluctuating conditions for two different species of juvenile coral reef fish, and other studies have found similar results (Jarrold and Munday 2018). Although fish in fluctuating future day conditions spent less time in the predator cue than fish in static future day conditions, their time spent was still higher than fish from both present day treatments, indicating that although the effect is reduced, it is still an issue of future concern. Given no significant differences between fluctuating and static conditions for present day treatments, fluctuations are more likely to play a crucial role in the future.

While our results support the overall conclusions of most behavioral studies investigating OA impacts on coral reef fishes (Munday et al. 2019), our results differ in the magnitude of these behavioral impairments. For example, the first chemosensory response study to report on predator detection found that under static future day conditions, settlement stage fish spent all of their time in predator cue, whereas those in static present day control conditions displayed a higher predator avoidance and spent no time in predator cue (Dixson et al. 2010). In contrast, when fish in our study were given the option between predator and non-predator cue, fish from the static future day treatment and static present day control spent 72.5% and 26.5% of time in predator cue, respectively. As the same focal species (*Amphiprion percula*), testing apparatus, and methods were used, a direct comparison can be made, potentially highlighting the role of juvenile age in response towards cues. Fish tested in Dixson et al. (2010) were treated during the egg and larval stage and tested at settlement (11 days post hatch), whereas the *A. percula* tested here were treated only during a two week period of the juvenile stage and tested at 20 weeks post hatch. Sensitivity to OA may likely be greater at earlier stages of ontogeny. As OA can affect larval processes, settlement and metamorphosis, early life stages represent a critical “bottleneck” period, meaning behaviors such as those tested in this study may be even more essential for survival at a younger age (Almany et al. 2006; Espinel-Velasco et al. 2018). Furthermore, our results suggest that fluctuations appear to provide little, if any, behavioral benefits under present day conditions. As coral reefs are generally not a homogenous environment, *p*CO_2_ ranges and fluctuations may vary both spatially across and within reefs, and temporarily, accounting for different behavioral and biological responses (Duarte et al. 2013; Boyd et al. 2016; Vargas et al. 2017).

As continual exposure to OA over time and movement from high to lower levels of *p*CO_2_ does not appear to reduce the negative behavioral impacts (Munday et al. 2014, 2016), fluctuating conditions and organismal adaptivity should be researched on a local scale, with a key emphasis on finetuning the different degrees of fluctuations and the important roles they play (Wahl et al. 2016). Given the variability and complexity of coral reefs, it remains largely unknown how *p*CO_2_ fluctuations may impact behavior, and on a larger scale, ecosystem structure and function (Queirós et al. 2014; Goldenburg et al. 2018).

## 5. Conclusion

The results found here underscore and expand on previous research that has assessed behavioral abnormalities of *Amphiprion percula* and other coral reef fishes under static future day *p*CO_2_ conditions (Munday et al. 2009, 2010; Dixson et al. 2010; Domenici et al. 2012; Ferrari et al. 2012a; Nilsson et al. 2012; Allan et al. 2013; Chivers et al. 2014). Furthermore, this study adds to the small yet growing literature suggesting that naturally occurring diel fluctuating *p*CO_2_ conditions may help mitigate or reduce OA-induced behavioral abnormalities under future climate change regimes.

## Acknowledgments

We thank the Dixson Lab for their assistance with experimental design and preparation for behavioral trials, D. Miller, J. Cohen, and P. Dominici for their assistance with statistical analysis, and the maintenance staff at the University of Delaware for their logistical assistance. All research was conducted under the guidelines of IACUC #1292. This research was funded by the National Science Foundation (Dixson #1750269). Data can be found at Zenodo (10.5281/zenodo.4459414)

